# Box C/D snoRNPs and MDT-15/MED15 regulate mitochondrial surveillance and mitophagy via fatty acid metabolism

**DOI:** 10.1101/2025.05.26.656193

**Authors:** Lois Armendariz, Alicia Chan, Elissa Tjahjono, Meggie Wang, Yvette Acevedo, Natalia V. Kirienko

**Affiliations:** Department of BioSciences, Rice University, 6100 Main St, MS140, Houston, TX, 77005, USA

## Abstract

In response to constant homeostatic threats, organisms have developed complex regulatory networks to monitor cellular functions and restore normal function. Here, we identify MDT-15 and its effectors, the fatty acid desaturases FAT-5, FAT-6, and FAT-7, as activators of the Ethanol and Stress Response (ESRE) mitochondrial surveillance pathway. Our data show that box C/D snoRNPs, which were previously linked to ESRE activation, also regulate FAT-6 and FAT-7 protein levels. Notably, knockdown of *mdt-15* or *fib-1*, a component of box C/D snoRNP complex, increased accumulation of the mitophagic activator PINK-1, the first step in licensing mitophagy, suggesting a relationship between ESRE surveillance and mitophagic activation. Supplementation with downstream unsaturated fatty acid products of FAT-6 and FAT-7 enhanced ESRE and mitophagic activation, but did not affect UPR^mt^. Since fatty acids activated ESRE and PINK-1 in wild-type and mutant genetic backgrounds, they are likely to act via a mechanism independent of FAT-6 and FAT-7 function. Our results provide insight into a novel interplay between box C/D snoRNPs, MDT-15, and fatty acids in the regulation of mitochondrial surveillance and mitophagy.

**Graphical Abstract:** 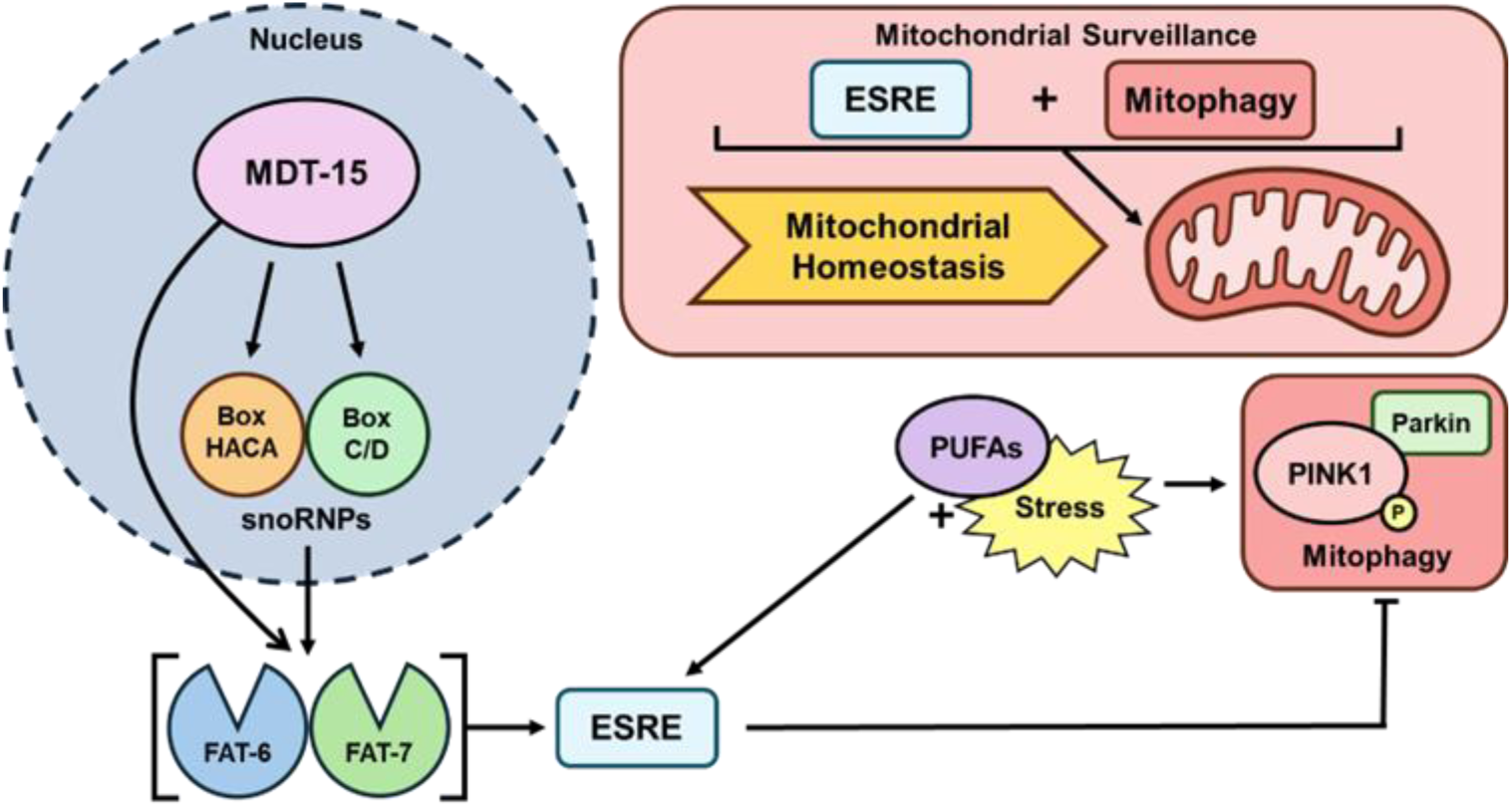

## Introduction

Mitochondria have long been characterized as “the powerhouse of the cell”. While there is some truth to this oversimplification, these organelles also play key roles in regulating a variety of important biochemical processes that determine cellular health. Compromising mitochondrial health can lead to metabolic and cardiovascular diseases, neurodegeneration, cancer, and aging (1). To maintain mitochondrial homeostasis, multicellular organisms have developed surveillance mechanisms and pathways that monitor a wide variety of signals, including mitochondrial bioenergetics, proteostasis, and key metabolites for disruptions that may indicate homeostatic imbalance. In response to these signals, the cell activates gene expression programs, coordinated between mitochondria and the nucleus, intended to restore normal cellular function (2).

The most studied of these pathways is the mitochondrial unfolded protein response (UPR^mt^) (3). This pathway activates the transcription factor ATFS-1, which upregulates mitochondrial chaperone effectors in response to mitochondrial dysfunction. Another, the PINK-1/Parkin mitophagic pathway, is responsible for autophagic degradation of mitochondria and selective respiratory chain turnover, which can be activated in basal or stressed conditions (4,5). A third mitochondrial surveillance system is the mitogen-activated protein kinase cascade (MAPK^mt^), defined by the DLK-1/SEK-3/PMK-3 proteins, which responds to mitochondrial bioenergetic abnormalities and plays a vital role in lifespan (6). Finally, the Ethanol and Stress Response Element (ESRE) surveillance pathway monitors reactive oxygen species (ROS), a common byproduct of mitochondrial function that plays an important role in aging (7). Kwon et al. first identified the ESRE motif as a conserved, 11-nucleotide DNA element (TCTGCGTCTCT) in the promoter region of genes upregulated in response to ethanol-induced stress (**Fig. 1A, Table S1**; note that all motifs shown in Fig. 1 were derived *de novo* from the gene lists in relevant publications using MEME) (8). In 2017, Tjahjono and Kirienko identified the same ESRE motif in genes upregulated in response to liquid-based pathogenesis with the Gram-negative opportunistic pathogen, *Pseudomonas aeruginosa* (LK-*Pa*), or acute iron removal, indicating an important role for ESRE in host defense (**Fig. 1B, Table S2**) (9). Follow-up research by Tjahjono et al. linked the ESRE pathway to mitochondrial surveillance, specifically to increased ROS from aberrations in the mitochondrial electron transport chain (7). This is consistent with other reports about the role of mitochondria in innate immunity (10). Activation of UPR^mt^ provides resistance to agar-based infections with *P. aeruginosa*, also known as Slow Killing (SK-*Pa*) (11,12).

**Figure 1.**
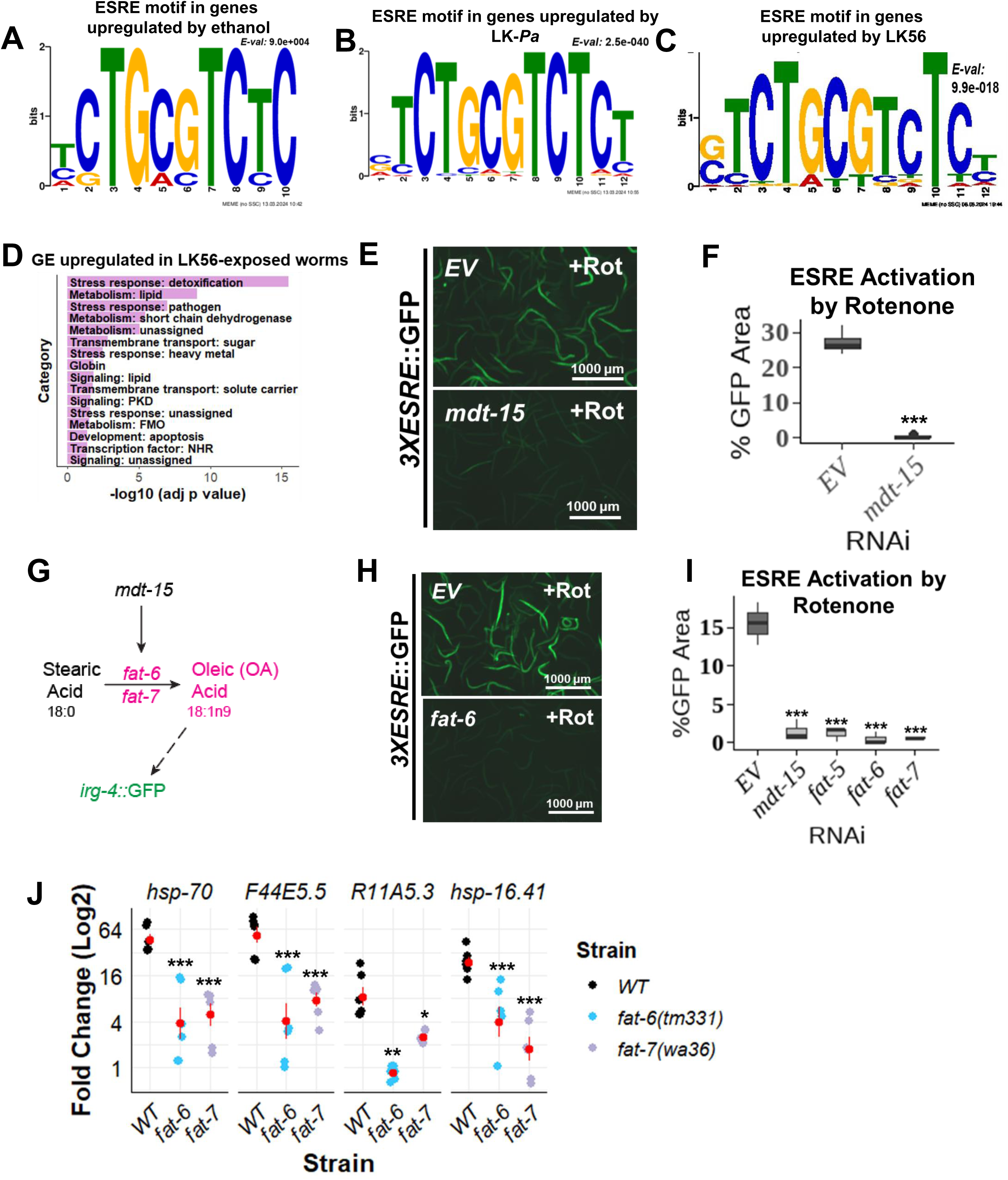
MDT-15 and its downstream effectors are required for activation of the Ethanol and Stress Response Element mitochondrial surveillance pathway. **(A-C)** *De novo* motifs identified in a subset of genes upregulated in response to 7% ethanol exposure **(A),** liquid-based *P. aeruginosa* pathogenesis **(B)**, or treatment with immune-stimulating compound, LK56 **(C)**. **(D)** Gene enrichment analysis (Wormcat) on genes upregulated in response to LK56 treatment. Horizontal bar graph depicts significantly enriched categories; adjusted *p* values were obtained by taking the negative logarithm base 10 of the original *p*-value. **(E,H)** Fluorescent images of *3XESRE*::GFP reporter strain reared on *EV(RNAi)*, *mdt-15(RNAi)* (E), *fat-5(RNAi), fat-6(RNAi)*, or *fat-7(RNAi)* (H) and treated with 25 μM rotenone. Quantification of percent GFP area, which gives a readout of ESRE activation **(F,I)**. **(G)** Diagram depicting a section of *C. elegans’* polyunsaturated fatty acid synthesis (PUFA) pathway where stearic acid (SA) is converted to oleic acid (OA) by the action of desaturases FAT-6/FAT-7, which are known to be regulated by MDT-15. It is also noted from previous research by Anderson et al. that OA is required for *irg-4*::GFP activation. **(J)** Rotenone-induced expression level of native ESRE genes in wild-type, *fat-6* mutant, or *fat-7* mutant worms. Gene expression levels were normalized to their DMSO-treated cohort. At least three biological replicates were performed for each experiment. Student’s t-test was performed to calculate the significance of a treatment or condition when there were two cohorts in the experimental setting. One-way analysis of variance (ANOVA) was performed to calculate the significance of a treatment or a condition when there were more than two cohorts in the experimental setting. No stars = not significant, *p < 0.05, ** p < 0.01, and *** p < 0.001.

Recently, there has been an increased appreciation for the diverse roles of lipid metabolism in cellular processes beyond structural integrity, storage, energy production, aging, and the regulation of proliferation (13,14). For example, monounsaturated fatty acids (MUFAs) like oleic acid play a role in innate immune activation in *Caenorhabditis elegans* (15). Additionally, lipids such as ceramides can provide protection from toxin- or pathogen-induced mitochondrial damage (16), further linking mitochondrial functions in energy metabolism and innate immune activation.

Another cellular component recently linked to immunometabolism is the box C/D small nucleolar ribonucleoprotein (snoRNP) complex. Canonically, snoRNPs modify ribosomal RNAs (rRNAs) or small nuclear RNAs (snRNAs) in a sequence-specific fashion, guided by small nucleolar non-coding RNAs (snoRNAs) (17,18). snoRNAs are divided into two groups, box C/D and box H/ACA, which canonically catalyze 2’-O-methylation and pseudouridylation modifications, respectively (19,20).

Each group of snoRNAs is associated with a different snoRNP complex, made up of non-overlapping groups of proteins. Several non-canonical roles for snoRNPs have been identified. For example, in response to doxorubicin-induced oxidative stress, Rpl13a (box C/D family) snoRNPs accumulate in the cytoplasm (21). Rpl13a snoRNPs also regulate the response to lipotoxic and oxidative stress (22–24). Box C/D and box H/ACA snoRNAs have been linked to innate immune activation, specifically by activating the protein kinase response (PKR), which can respond to dsRNA or metabolic stress (23,25). Finally, previous research in our lab identified a role for box C/D snoRNPs in the regulation of mitochondrial surveillance, with the most prominent effect on activation of ESRE (26).

In this study, we identify roles for the evolutionarily-conserved mediator subunit MDT-15/MED15, the fatty acid desaturases FAT-5, FAT-6, and FAT-7, fatty acids, and box C/D snoRNPs in the regulation of mitochondrial surveillance and mitophagy. MDT-15, a major regulator of lipid metabolism, and its downstream effectors FAT-5, FAT-6, and FAT-7, were required for mitochondrial ESRE and UPR^mt^ activation. In addition, we uncover a non-canonical role for box C/D snoRNPs in the regulation of fatty acid metabolism through a FAT-6/FAT-7 axis, and that MDT-15 is required for basal snoRNP expression. Furthermore, we show increased mitophagic activation in response to suppressed ESRE. Additionally, we demonstrate that fatty acids from *C. elegans’* polyunsaturated fatty acid (PUFA) synthesis pathway can enhance activation of ESRE and mitophagy, but not UPR^mt^. Taken together, we elucidate a novel relationship between ESRE, box C/D snoRNPs, MDT-15, mitophagy, and fatty acids.

## RESULTS

### The evolutionarily conserved mediator subunit MDT-15/MED15 is required for ESRE activation

While investigating the biology underlying the host response to *P. aeruginosa* in Liquid Killing (LK), we carried out a high-throughput screen to identify small molecules that increase *C. elegans* survival (27–29). Five hits stimulated host innate immunity, with LK56 providing the most protection (28). Although transcriptional profiling was performed on LK56-treated worms and the compound was shown to depend on MDT-15 for its protective activity, a mechanism of action was not identified.

To uncover the potential mechanism of LK56-mediated rescue, upstream regulatory sequences from genes upregulated by LK56 were analyzed using RSAT (30) and MEME (31). Analyses revealed significant enrichment of the ESRE motif (30.7% of genes contained the ESRE motif) in their promoter regions **(Fig. 1C, Table S3)**. Analysis of LK56-upregulated genes using WormCat (32) found lipid metabolism, detoxification, and pathogen response as the most enriched categories **(Fig. 1D)**. This was consistent with previous research from the Kirienko lab showing that LK56 increased host resistance to *P. aeruginosa, Enterococcus faecalis,* and *Staphylococcus aureus* (28). Analysis of ESRE genes upregulated by LK56 using WormCat (32) also found similar enrichment in the categories of lipid metabolism and detoxification stress response **(Fig. S1A)**. Since the ESRE motif was enriched in the promoter regions of LK56-responsive genes and LK56 activity depends on MDT-15, we hypothesized that activation of ESRE genes may similarly rely on MDT-15. To test this, RNA interference (RNAi) was used to knock down *mdt-15* in a *C. elegans* strain carrying a GFP reporter driven by three tandem repeats of the ESRE motif (*3XESRE*::GFP) (33). This reporter strain was exposed to rotenone, which is known to activate ESRE (7). *mdt-15(RNAi)* virtually abolished GFP expression, even in the presence of rotenone **(Fig. 1E-F)**. These results indicate that the evolutionarily-conserved mediator subunit MDT-15 is required for ESRE activation in response to mitochondrial damage.

### FAT-5, FAT-6, and FAT-7 link ESRE activation to lipid metabolism

Amongst its many targets, MDT-15 has previously been shown to regulate the expression of the *Δ9*-fatty acid desaturases FAT-5, FAT-6, and FAT-7 (34). FAT-5 converts palmitic acid into palmitoleic acid (PA), while FAT-6 and FAT-7 convert the saturated fat stearic acid (SA) into monounsaturated oleic acid (OA) **(Fig. 1G)**. Desaturation of SA is required for proper membrane function, and MUFAs play key roles in lipid distribution, fat storage, and signaling (35). We tested the effect of *fat-5(RNAi)*, *fat-6(RNAi),* or *fat-7(RNAi)* on rotenone-mediated *3XESRE*::GFP activation **(Fig. 1H-I)**. Like MDT-15, these effectors were required for full activation of the ESRE reporter. Importantly, genetic mutants *fat-6(tm331)* and *fat-7(wa36)* showed significantly reduced expression of native ESRE genes in qRT-PCR **(Fig. 1J)**, confirming that MUFA production is necessary for activation of this mitochondrial surveillance pathway.

### FAT-6 and FAT-7 are activated in response to mitochondrial damage in a box C/D snoRNP-dependent manner

Previously, Tjahjono et al. described a non-canonical role for box C/D small nucleolar ribonucleoproteins (snoRNPs) in mitochondrial surveillance and innate immunity. The box C/D snoRNP protein complex, comprised of FIB-1, NOL-56, and NOL-58, act as a molecular switch that activates mitochondrial surveillance (ESRE and UPR^mt^) while repressing the PMK-1/p38 MAPK innate immune pathway (26). For this reason, we analyzed the relationship between box C/D snoRNPs and fatty acid metabolism genes in ESRE regulation using reporters carrying a *fat-7p*::FAT-7::GFP or a *fat-*6*p*::FAT-6::GFP array (36,37). The effect of *mdt-15(RNAi)* or *fib-1(RNAi)* on FAT-7::GFP levels during mitochondrial stress induced by the iron chelator, 1,10-phenanthroline (38,39) was studied **(Fig. 2A-B)**. Our results indicate that FAT-7 protein levels increase during mitochondrial stress in an MDT-15- and box C/D snoRNP-dependent manner.

**Figure 2.**
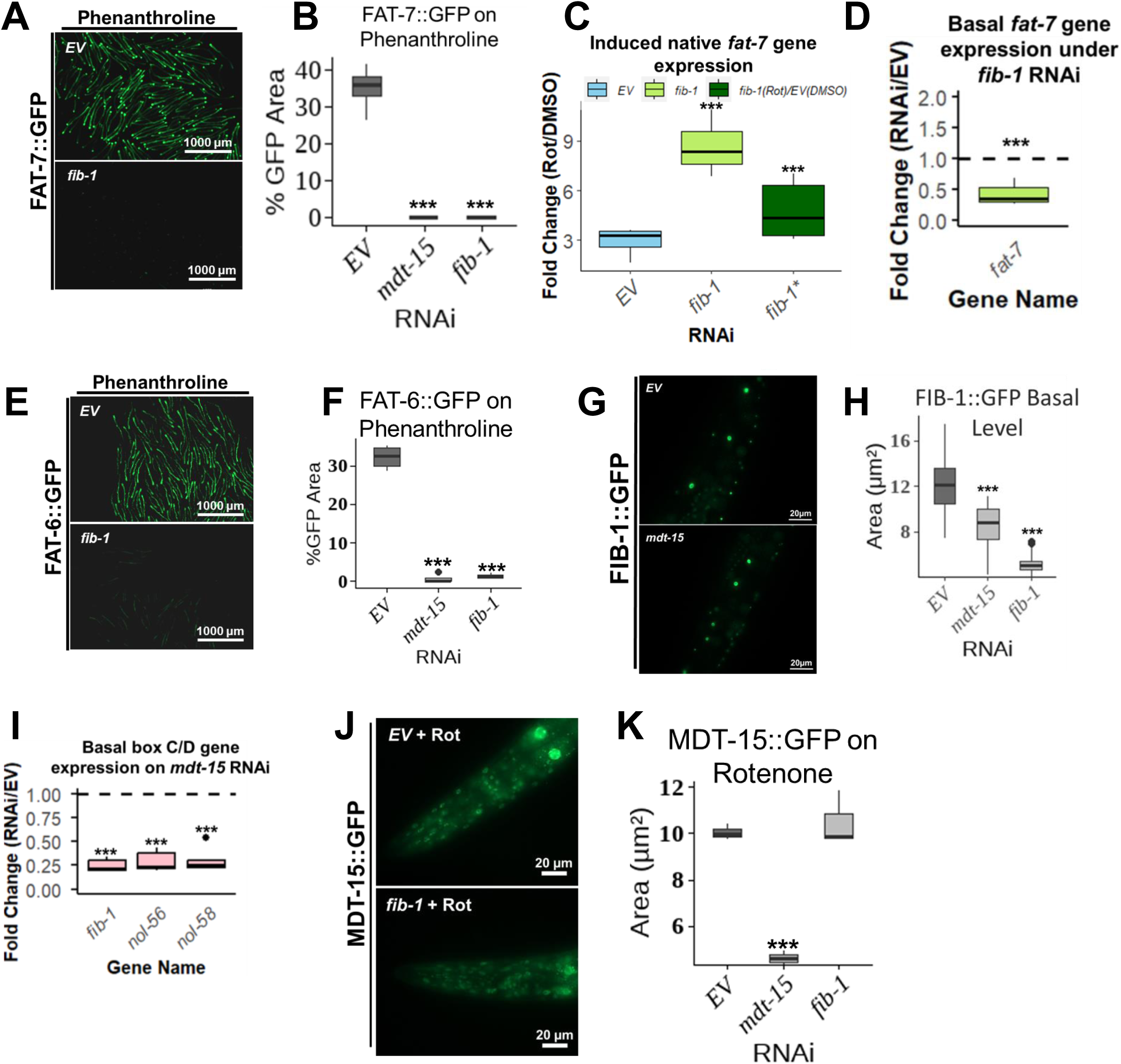
MDT-15, FAT-6, FAT-7, and box C/D snoRNPs coordinate to regulate ESRE. **(A,E)** Fluorescent images of FAT-7::GFP **(A)** or FAT-6::GFP **(E)** worms reared on *fib-1(RNAi)* and treated with phenanthroline. **(B,F)** Quantification of percent GFP area, which gives a readout of FAT-7 **(B)** or FAT-6 **(F)** levels. **(C,D)** Expression levels of native *fat-7* expression in worms reared on *EV* or *fib-1* RNAis and treated with 25 μM rotenone or *v/v* DMSO. Rotenone-induced gene expression levels were either normalized to their DMSO cohort or to *EV(RNAi)* DMSO levels **(C)**, or *fib-1(RNAi)* DMSO levels were normalized to *EV(RNAi)* DMSO levels **(D)**. **(G)** Fluorescent images of FIB-1::GFP worms reared on *EV* or *mdt-15* RNAis. **(H)** Area of FIB-1 punctae in FIB-1::GFP worms reared on *EV*, *mdt-15*, or *fib-1* RNAis and treated with DMSO. **(I)** Native gene expression of box C/D snoRNPs of worms fed with *EV(RNAi)* or *mdt-15(RNAi)*. Gene expression levels were normalized to the negative RNAi *(EV)* control. **(J)** Fluorescent images of MDT-15 punctae in worms reared on *EV* or *fib-1* RNAis and treated with 25 μM rotenone. **(K)** Area of MDT-15 punctae in MDT-15::GFP worms reared on *EV*, *mdt-15*, or *fib-1* RNAis and treated with rotenone. One-way analysis of variance (ANOVA) was performed to calculate the significance of a treatment or a condition when there were more than two cohorts in the experimental setting. No stars = not significant, *p < 0.05, ** p < 0.01, and *** p < 0.001.

To gain insight into the regulatory mechanism used by box C/D snoRNPs, qRT-PCR was performed to evaluate *fat-7* expression under *fib-1(RNAi)* in ESRE-activating conditions. In contrast to translational reporter results, *fat-7* expression upon rotenone exposure was upregulated in *fib-1(RNAi)* worms **(Fig. 2C)**. Since the basal level of native *fat-7* expression was lower under *fib-1(RNAi)* (**Fig. 2D**), we also normalized the rotenone-induced *fat-7* level in *fib-1(RNAi)* to the basal *EV(RNAi) fat-7* level and still observed significant upregulation (**Fig 2C**, dark green). The difference between transcriptional and translational results suggests that box C/D snoRNPs regulate FAT-7 post-transcriptionally (see Discussion). Similar results were observed with phenanthroline-induced increase in FAT-6::GFP levels **(Fig. 2 E-F)**.

### Expression of FIB-1 depends on MDT-15

To uncover the connection between box C/D snoRNPs and fatty acid metabolism, a worm strain carrying a single-copy insertion of FIB-1::GFP driven by its native promoter (40) was used. Worms were reared on RNAi targeting *mdt-15* or *fib-1,* and the effects on basal *fib-1* expression were evaluated **(Fig. 2 G-H)**. *mdt-15(RNAi)* reduced the area of FIB-1 punctae, suggesting an unanticipated regulatory mechanism wherein MDT-15 drives expression of box C/D snoRNPs.

The effect of *mdt-15(RNAi)* on native box C/D gene expression was also evaluated. *mdt-15(RNAi)* reduced expression of all genes tested, as compared to the negative RNAi (*EV)* control, supporting the conclusion that MDT-15 transcriptionally regulates box C/D snoRNPs **(Fig. 2I).** To test whether box C/D snoRNPs reciprocally regulate MDT-15, an MDT-15::GFP reporter strain was used (41). Worms were reared on RNAi-inducing *E. coli* targeting *mdt-15* or *fib-1* and then treated with rotenone. **(Fig. 2 J-K).** Knockdown of *fib-1* did not affect levels of MDT-15 protein, indicating that MDT-15 is at the top of this signal transduction network.

#### Suppression of ESRE surveillance results in enhanced mitophagic activation

Another notable mitochondrial surveillance pathway is the PINK-1/Parkin pathway, which controls selective autophagic degradation of mitochondria (mitophagy) (4,5). To assess the relationship between mitophagy and ESRE, a *pink-1p*::PINK-1::GFP reporter was used (42). Worms were reared on RNAi targeting *mdt-15, fat-6,* or *fib-1*, and then were treated with *trans*-β-nitrostyrene (nitrostyrene, **Fig. 3 A-C)**, a robust mitophaic activator in *C. elegans* and human cells (42,43). Knockdown of *mdt-15* or *fib-1* significantly increased PINK-1 accumulation and mitophagic activation. To verify that the increase in PINK-1 levels under these conditions leads to autophagic activation, a strain carrying a *lgg-1p*::LGG-1::GFP translational fusion was used (44,45). Knockdown of *fib-1* increased LGG-1::GFP punctae upon treatment with nitrostyrene, while a double *fib-1(RNAi); pink-1(RNAi)* knockdown decreased LGG-1::GFP punctae, showing that *fib-1* increases autophagy in a PINK-1-dependent manner **(Fig. 3 D-E)**. As FIB-1 and MDT-15 are required for ESRE activation, reduced ESRE activity is likely to increase mitophagy, we predict as a compensatory mechanism. Similarly, this would suggest that activated ESRE may limit mitophagy by improving mitochondrial function, revealing a novel co-regulatory relationship between mitochondrial surveillance and mitophagy.

**Figure 3.**
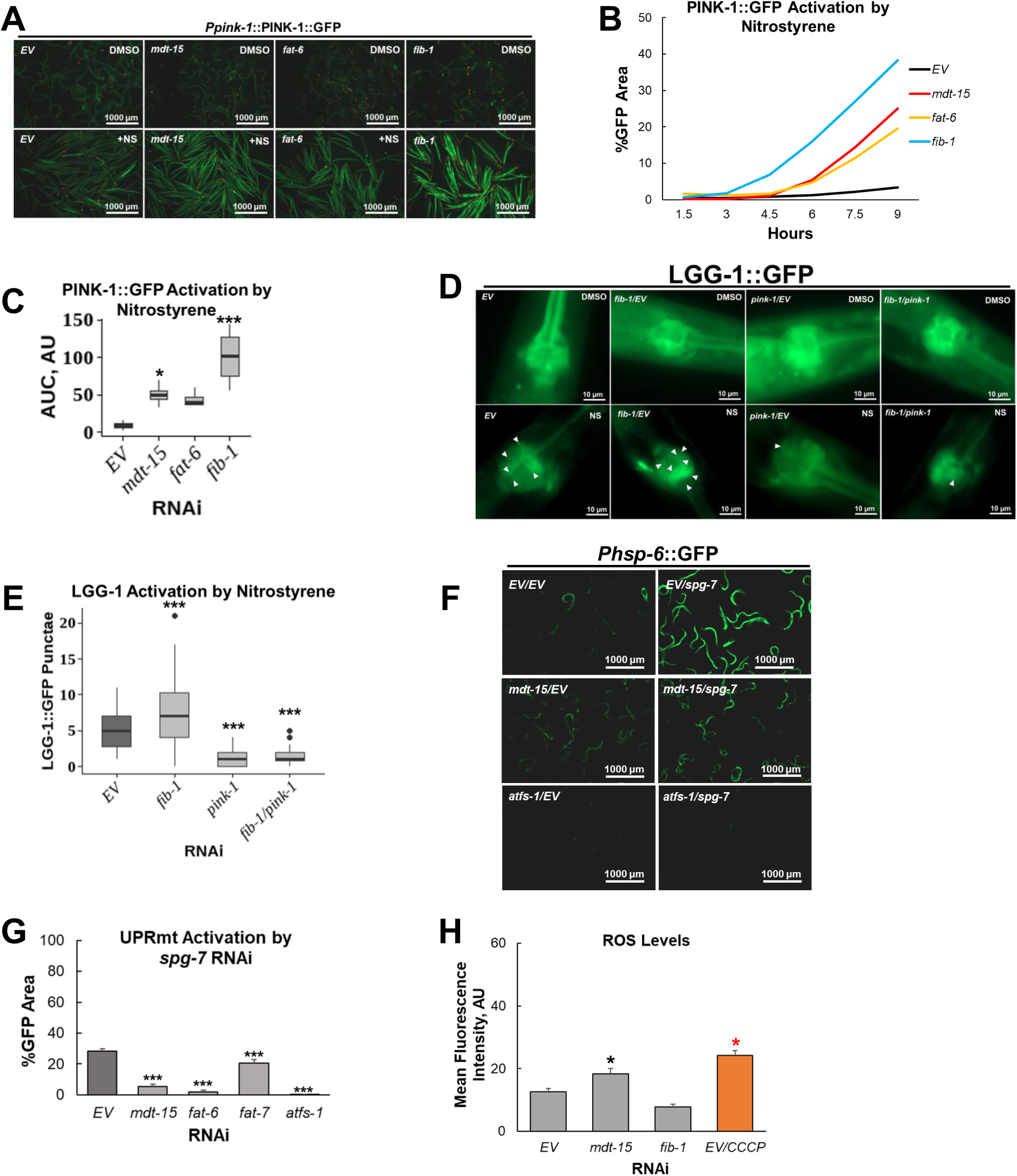
ESRE regulators affect PINK-1 and UPR^mt^ activation. **(A)** Fluorescent images of PINK-1::GFP worms reared on *EV*, *mdt-15*, *fat-6*, or *fib-1* RNAis and treated with 30 µM nitrostyrene or v/v DMSO. **B)** Curve representing PINK-1::GFP activation by nitrostyrene across various hours. **(C)** Area under the curve quantification for PINK-1::GFP activation by nitrostyrene across various hours. **(D)** Fluorescent images of LGG-1::GFP worms reared on *EV*, *pink-1*, or *fib-1* RNAis and treated with 30 μM nitrostyrene or *v/v* DMSO. White arrows point to LGG-1 punctae in the worm’s pharyngeal terminal bulb. **(E)** Quantification of LGG-1 punctae for each condition. **(F)** Fluorescent images of *Phsp-6*::GFP worms reared on *EV*, *mdt-15,* or *atfs-1* RNAis. Each RNAi was mixed in a 3:1 ratio with *spg-7* or *EV* RNAis. **(G)** Quantification of percent GFP area, which gives a readout of UPR^mt^ activation. **(H)** Quantification of DCFDA mean fluorescence intensity, which gives a readout of ROS levels. *glp-4(bn2)* worms were reared on *EV, mdt-15,* or *fib-1* RNAis and treated with either 78.2 μM CCCP or *v/v* DMSO.

#### MDT-15 regulates other mitochondrial surveillance pathways

Multicellular organisms have numerous mitochondrial surveillance pathways that respond to a variety of stresses (2). We tested whether MDT-15, FAT-6, or FAT-7 also regulate the mitochondrial unfolded protein response (UPR^mt^). RNAi was used to knock down *mdt-15*, *fat-6*, or *fat-7* in a reporter strain carrying *hsp-6p*::GFP, a reporter for UPR^mt^ (46), which was activated by *spg-7(RNAi)* **(Fig. 3 F-G)**. Knocking down *mdt-15*, *fat-6*, or *fat-7* compromised activation of UPR^mt^.

Since ROS are critical for ESRE induction, we tested whether FIB-1 or MDT-15 activate ESRE by increasing ROS levels. For this, *glp-4(bn2)* worms were reared on RNAi targeting *mdt-15* or *fib-1* and then stained with 2’,7’ -dichlorofluorescin diacetate (DCFDA). We confirmed function of the assay by treating worms reared on a control RNAi with carbonyl cyanide *m-*chlorophenyl hydrazone (CCCP), which is known to induce ROS (7). CCCP is an ionophore that inhibits mitochondrial oxidative phosphorylation, inducing mitochondrial dysfunction and oxidative stress via ROS production (47). Our results show that *mdt-15*(*RNAi*) raised basal ROS levels, while *fib-1(RNAi)* had no effect **(Fig. 3H)**, but none of the knockdowns resulted in ROS depletion. This indicates that FIB-1 and MDT-15 are more likely to be directly involved in the signal transduction required for ESRE activation rather than indirectly regulating ESRE by impacting ROS.

### Exogenous supplementation of oleic acid enhances ESRE activation

FAT-6 and FAT-7 are known to be involved in OA production **(Fig. 4A)** and innate immune activation (15). To determine whether OA level affects ESRE activation, RNAi was used to knock down *mdt-15*, *fat-6*, or *fat-7* in worms carrying the *3XESRE*::GFP reporter reared on media supplemented with 500 μM OA **(Fig. 4 B-D, compare dashed to solid lines and blue to gold boxplots per RNAi)**. The addition of OA enhanced rotenone-triggered ESRE activation in *fat-6* or *fat-7*, but not *mdt-15* knockdowns (red stars indicate significant AUC levels between blue and gold boxplots per RNAi). Similar results were seen regardless of whether the non-ionic surfactant Tergitol (NP-40) was added to the medium (compare **Fig. S1 B-C** and **Fig. 4 B-D)**. Since the assay was working well in the absence of Tergitol, subsequent experiments involving supplementation were done using DMSO as the vehicle). Interestingly, OA supplementation also augmented rotenone-triggered ESRE activation in worms reared on the empty vector (*EV*) negative RNAi control, suggesting that OA limitation normally curtails ESRE mitochondrial surveillance. However, OA alone was insufficient to induce ESRE pathway activation in the absence of mitochondrial stress triggered by rotenone **(Fig. S1 D-G)**. This indicates that OA augments activation under stress, but does not cause mitochondrial damage on its own.

**Figure 4.**
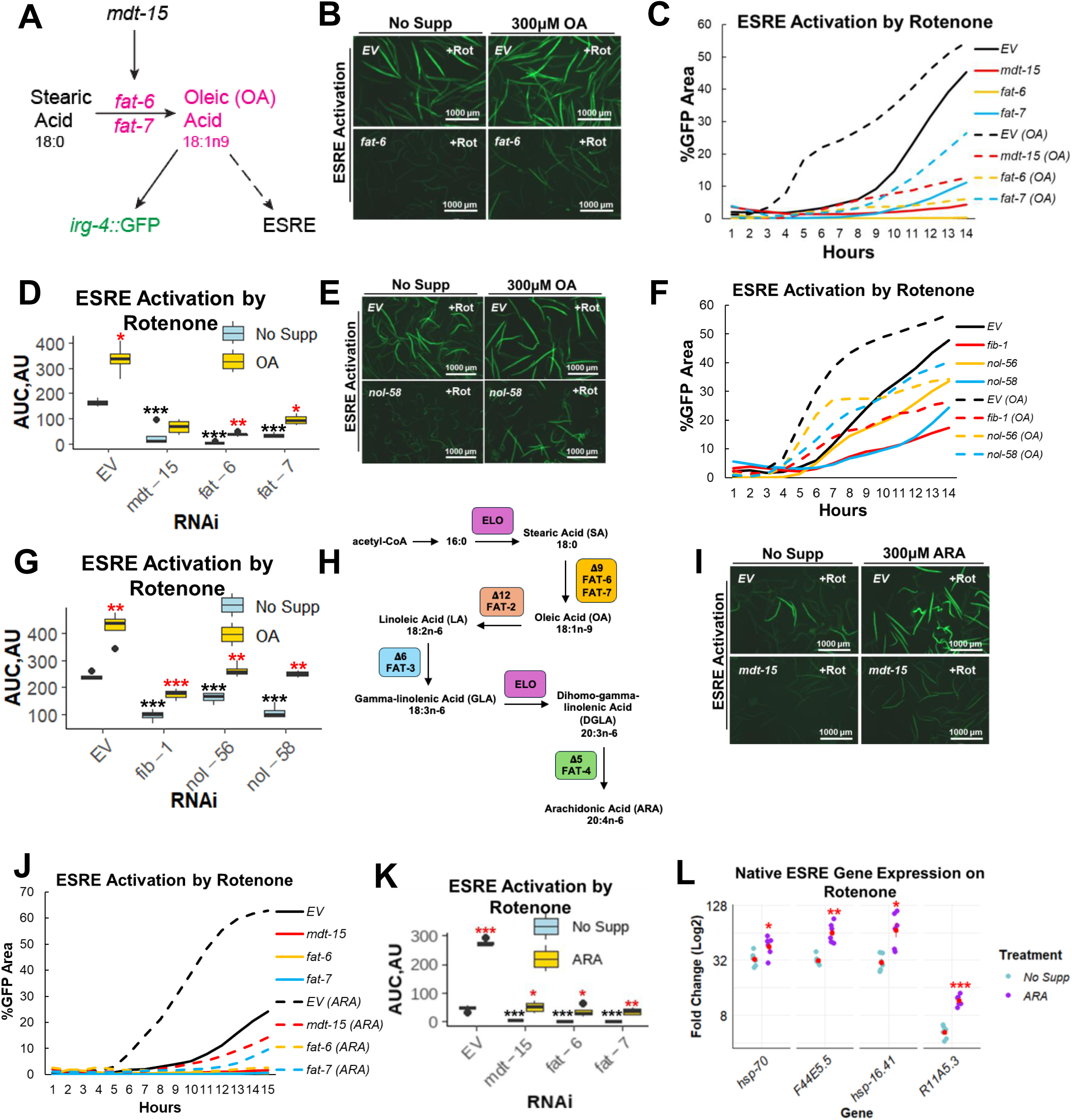
The mediator subunit MDT-15 and its downstream effectors *fat-6* and *fat-7* are required for ESRE activation. **(A)** Diagram depicting a section of *C. elegans’* PUFA pathway where stearic acid (SA) is converted to oleic acid (OA) by the action of desaturases FAT-6/FAT-7, which are known to be regulated by MDT-15. It is also noted from previous research by Anderson et al. that OA is required for *irg-4*::GFP activation. The diagram also shows that OA is required for ESRE activation. **(B,I)** Fluorescent images of *3XESRE*::GFP worms reared on *EV(RNAi), fat-6(RNAi), mdt-15(RNAi),* or *nol-58(RNAI)* **(E)** supplemented with OA **(B, E)** or ARA **(I)** and treated with 25 μM rotenone. **(C,F,J)** Curve representing ESRE activation by rotenone across various hours. **(D,G,K)** Area under the curve quantification for ESRE activation by rotenone across various hours. **(H)** Diagram depicting *C. elegans’* PUFA pathway. **(L)** Rotenone-induced expression level of native ESRE genes in wild-type worms supplemented/not supplemented with ARA. Gene expression levels were normalized to their DMSO-treated cohort. In all panels, red stars indicate comparisons between fatty acid supplementation and no supplementation for each RNAi or gene. In all panels, black stars indicate comparisons between RNAi knockdowns and *EV(RNAi)* negative control in the non-supplemented cohort. Student’s t-test was performed to calculate the significance of a treatment or condition when there were only two cohorts in the experimental setting. One-way analysis of variance (ANOVA) was performed to calculate the significance of a treatment or a condition when there were more than two cohorts in the experimental setting. No stars = not significant, *p < 0.05, ** p < 0.01, and *** p < 0.001.

Since OA enhanced ESRE activation in *fat-6* and *fat-7* knockdowns, and box C/D snoRNPs are involved in ESRE activation, we tested the effect of exogenous OA supplementation on worms with knocked down snoRNP complex proteins. Worms carrying the ESRE reporter were fed RNAi targeting *fib-1*, *nol-56*, or *nol-58*, supplemented with OA, and treated with rotenone. Rotenone-triggered ESRE expression was enhanced by OA, but expression was not restored to normal levels. The same results were observed when *fat-6(RNAi)* and *fat-7(RNAi)* were tested under the same conditions **(Fig. 4 E-G, compare blue to gold boxplot per RNAi)**. These data suggest that OA may function in parallel to box C/D snoRNPs in ESRE activation.

### Unsaturated fatty acid supplementation enhances activation of ESRE and mitophagy, but not UPR^mt^

Rotenone-treated *EV(RNAi)*, *fat-6(RNAi), fat-7(RNAi),* and *fib-1(RNAi)* each showed similar OA-mediated increases in ESRE reporter expression. This suggests that OA functions in parallel to these genes. To examine the role of downstream fatty acids in the *C. elegans* PUFA pathway **(Fig. 4H)** in mitochondrial surveillance, knockdowns of *mdt-15, fat-6*, *fat-7,* or *fib-1* in the *3XESRE*::GFP background were reared on media containing 500 μM linoleic acid (LA), γ-linolenic acid (GLA), or arachidonic acid (ARA). Supplementation with any of these fatty acids enhanced ESRE expression after rotenone exposure **(Fig. 4 I-K, Fig. S2 A-I, compare blue to gold boxplot per RNAi; compare blue to orange, green, or pink boxplot for *fib-1* RNAi)**. Importantly, the addition of ARA also increased expression of native ESRE genes after rotenone exposure **(Fig. 4L, compare purple to blue dots per gene)**, demonstrating that this role is biologically relevant endogenous ESRE gene activation.

Interestingly, when *mdt-15(RNAi);* FIB-1::GFP worms were reared on ARA-supplemented media, we saw no increase in the size of FIB-1 punctae compared to the negative RNAi control (*EV)*. This suggests that ESRE regulation by fatty acids is regulated independently of the MDT-15/FIB-1 axis **(Fig. S2I, compare magenta and pink to blue per RNAi)**.

Next, we tested whether fatty acid supplementation affected other mitochondrial surveillance pathways. Supplementation with ARA further increased PINK-1::GFP reporter expression in the *EV* control or in *mdt-15*, *fat-6*, or *fib-1* RNAi knockdowns, similar to ESRE (**Fig**. **5 A-B****, Fig. S3A compare blue to gold boxplots per RNAi)**. We also tested whether fatty acid supplementation could enhance activation of the UPR^mt^ reporter *hsp-6p*::GFP after *mdt-15*, *fat-6,* or *fat-7* were knocked down. In contrast to ESRE, neither OA, LA, nor ARA supplementation increased *spg-7(RNAi)*-mediated expression of the reporter **(Fig. 5C, S3 B-F compare gold to blue bar graphs per RNAi)**. Instead, OA decreased UPR^mt^ activation in *fat-7(RNAi)* and the negative RNAi (*EV)* control, LA decreased activation in the negative RNAi (*EV)* control, and ARA supplementation decreased activation in *fat-6(RNAi)*. These data indicate that fatty acid-mediated enhancement of mitochondrial surveillance is specific to ESRE and mitophagy.

**Figure 5.**
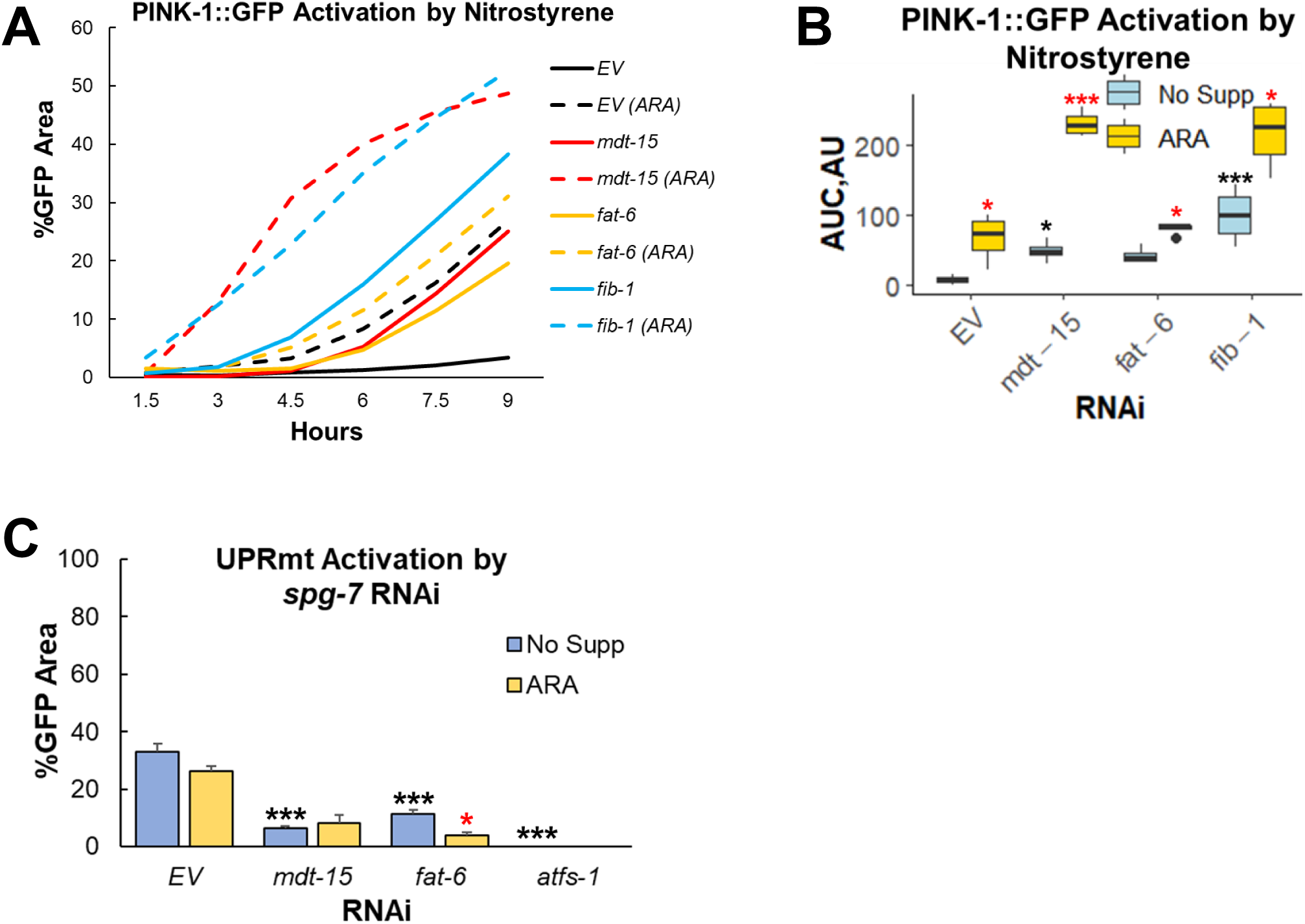
Fatty acid supplementation enhances PINK-1 activation but not UPR^mt^. **(A)** Curve representing PINK-1::GFP activation by nitrostyrene across various hours. **(B)** Area under the curve quantification for PINK-1::GFP activation by rotenone across various hours. **(C)** Quantification of percent GFP area, which gives a readout of UPR^mt^ activation. In all panels, red stars indicate comparisons between fatty acid supplementation and no supplementation for each RNAi. In all panels, black stars indicate comparisons between RNAi knockdowns and *EV(RNAi)* negative control in the non-supplemented cohort. Student’s t-test was performed to calculate the significance of a treatment or condition when there were only two cohorts in the experimental setting. One-way analysis of variance (ANOVA) was performed to calculate the significance of a treatment or a condition when there were more than two cohorts in the experimental setting. No stars = not significant, *p < 0.05, ** p < 0.01, and *** p < 0.001.

### Supplementation of fatty acids during mitochondrial stress increases oxidative stress

We hypothesized that fatty acids may enhance ESRE activation by increasing ROS levels during rotenone-induced mitochondrial stress. To test this, we treated worms with ascorbate (vitamin C), a known antioxidant (48) that was shown to inhibit ROS-induced ESERE activation. Ascorbate abrogated ESRE activation in rotenone-treated worms regardless of ARA supplementation, likely by scavenging ROS **(Fig. 6 A-B**, **compare blue to gold boxplots per treatment).** This is consistent with our previous findings that rotenone induces ESRE activation by generating ROS (7) and indicates that fatty acids may enhance ESRE activation by increasing ROS. In parallel, we measured basal ROS levels of worms reared in media supplemented with OA, LA, GLA, or ARA **(Fig. 6C)** and saw no significant change from non-supplemented worms, strengthening our hypothesis that fatty acids do not cause stress by themselves in the absence of mitochondrial stress.

**Figure 6.**
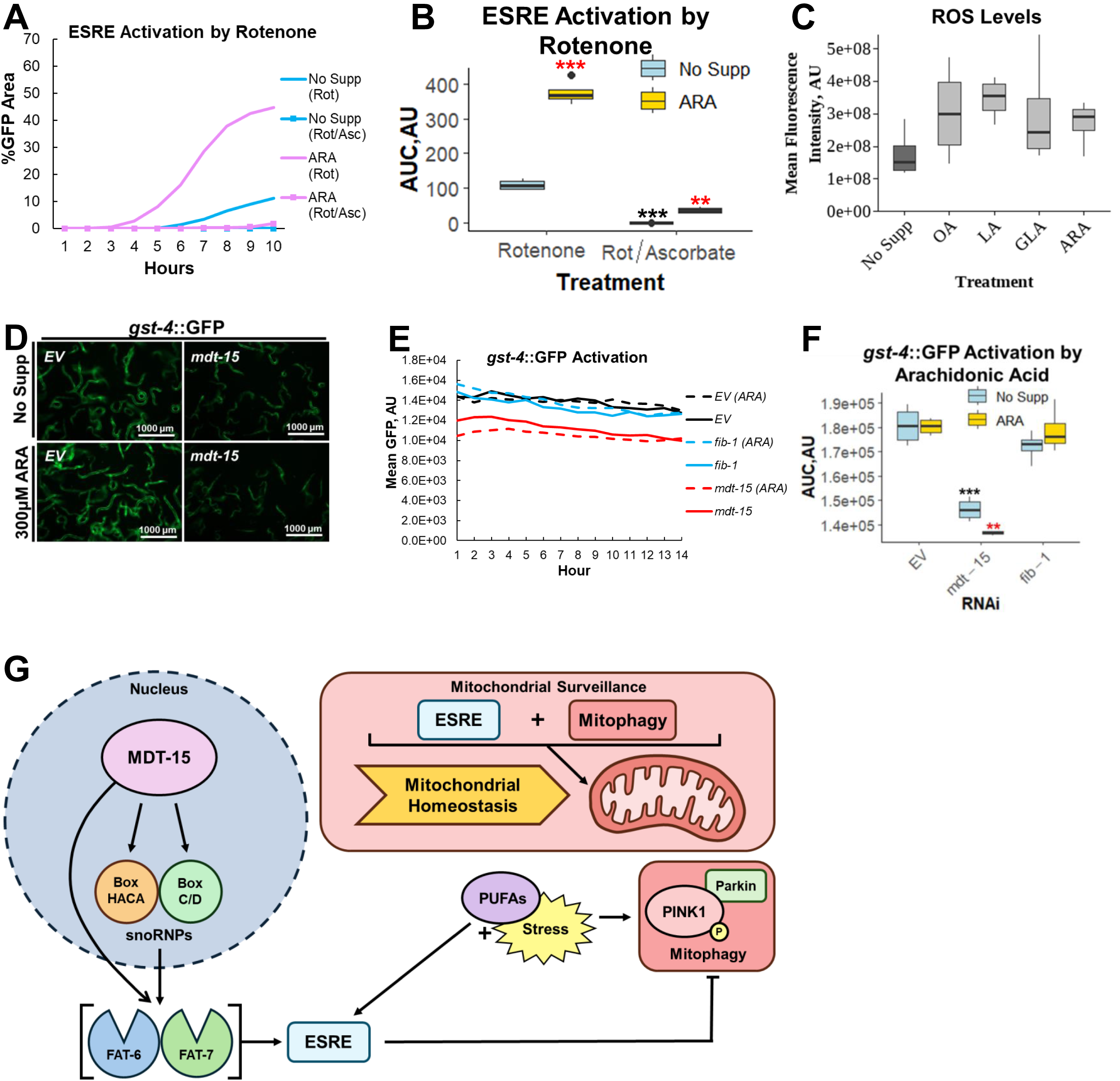
Fatty acid supplementation by itself does not induce oxidative stress. **(A)** Curve representing ESRE activation by rotenone across various hours. **(B)** Area under the curve quantification for ESRE activation by rotenone across various hours. **(C)** Quantification of DCFDA mean fluorescence intensity, which gives a readout of ROS levels. *glp-4(bn2)* worms were reared on media supplemented with OA, LA, GLA, or ARA. **(D)** Fluorescent images of *gst-4*::GFP worms reared on *EV* or *mdt-15* RNAis supplemented with ARA. **(E)** Curve representing *gst-4*::GFP activation by ARA across various hours. **(F)** Area under the curve quantification for *gst-4*::GFP activation by ARA across various hours. **(G)** Model depicting the pathways by which MDT-15 regulates box C/D snoRNPs, mitochondrial surveillance, and mitophagy. In all panels, red stars indicate comparisons between fatty acid supplementation and no supplementation for each RNAi. In all panels, black stars indicate comparisons between RNAi knockdowns and *EV(RNAi)* negative control in the non-supplemented cohort. Student’s t-test was performed to calculate the significance of a treatment or condition when there were only two cohorts in the experimental setting. One-way analysis of variance (ANOVA) was performed to calculate the significance of a treatment or a condition when there were more than two cohorts in the experimental setting. No stars = not significant, *p < 0.05, ** p < 0.01, and *** p < 0.001.

We also tested the impact of fatty acid supplementation on activation of the oxidative stress response using a *gst-4*::GFP reporter (49). RNAi knockdowns of *mdt-15* or *fib-1* in the *gst-4*::GFP background were reared on media containing 500 μM ARA **(Fig. 6 D-F)**. Supplementation of ARA did not trigger oxidative stress by itself, validating our basal ROS measurement results.

## Discussion

In this study, we found that MDT-15/MED15 and box C/D snoRNPs work together to modulate mitochondrial surveillance and mitophagy, likely by regulating fatty acid metabolism **(Fig. 6G)**. This recapitulates and extends earlier findings, where *mdt-15*(*-*) mutants showed differences in expression of a number of small HSPs, including *hsp-16.1, hsp-16.41, hsp-16.2,* and *F44E5.5* (41). They observed that the *mdt-15*(-) mutant had an altered ratio of unsaturated to saturated fatty acids due to differential expression of *fat-6* and *fat-7*. They also noted that supplementation with OA restored many of the *mdt-15*(*-*) phenotypes, and they connected these phenotypes to membrane fluidity, possibly sensed by PAQR-2.

Although our paper agrees with that report in most respects, there is a significant difference worth discussing. Lee et al. concluded that the proteostatic effects they observed were independent of mitochondrial stress, as *hsp-6*, *hsp-60*, and *Y22D7AL.10* expression levels were all low in *mdt-15*(-) mutants. They also observed increased cytosolic chaperone expression with decreased fat levels, which is inconsistent with the mitochondrial-to-cytosolic stress response (mitoCSR) (50), furthering their confidence that mitochondrial stress was not involved in the response observed.

However, the chaperones whose gene expression profiles they measured are associated with UPR^mt^, but not with the ESRE mitochondrial surveillance pathway. Interestingly, several of the chaperones that were differentially regulated by *mdt-15*(*-*) mutants at low temperatures, including *hsp-16.1, hsp-16.2, hsp-16.11, hsp-16.41, hsp-16.49, F44E5.4,* and *F44E5.5,* have easily recognized ESRE elements in their promoter regions, and *hsp-16.1, hsp-16.41*, and *atfs-1* have been used to demonstrate the importance of the ESRE site in appropriate context-dependent gene expression (7,51). This paper demonstrates that the phenomenon observed by Lee, et al. may include an unrecognized mitochondrial component, albeit one controlled by ESRE rather than the UPR^mt^, which only partially overlap (7,26)this paper).

Using bioinformatics approaches (i.e., Wormcat, RSAT, and MEME SUITE) to investigate the immune-stimulating compound LK56, we found enrichment of upregulated lipid metabolism genes, several of which contained ESRE motifs in their promoters (30–32). Similar results were observed when we looked at genes upregulated in response to *P. aeruginosa* liquid-based pathogenesis, indicating strong connections between lipid metabolism, host defense, and mitochondrial surveillance. Using reporter strains and gene expression analyses of native ESRE genes, we demonstrated the involvement of MDT-15, FAT-6, and FAT-7 in the regulation of the mitochondrial ESRE and UPR^mt^ pathways, strengthening the link between lipid metabolism and the regulation of mitochondrial surveillance. This was an interesting finding, as the role of lipids like ceramide in the regulation of mitophagy and activation of UPR^mt^ during proteotoxic stress has been established (16,52,53), but a role for lipids in ESRE regulation is novel.

Since MDT-15, FAT-6, and FAT-7 are required for ESRE and UPR^mt^, we hypothesized that exogenous supplementation of downstream fatty acids would rescue their activation if *mdt-15, fat-6,* or *fat-7* were disrupted by RNAi. To our surprise, fatty acid supplementation augmented ESRE activation in wild-type and mutant backgrounds, suggesting a separate regulatory mechanism, rather than a specific rescue. Interestingly, fatty acid supplementation without additional mitochondrial stress did not activate ESRE or mitophagy pathways. Instead, supplementation specifically enhanced the activation of these pathways under stressed conditions. In contrast, no augmentation by fatty acid supplementation was detected for the UPR^mt^. The likeliest explanation for this difference is the greater sensitivity the ESRE response has for ROS than the UPR^mt^ does (7); increased levels of long-chain fatty acids can impair mitochondrial redox activity and enhance steady-state production of ROS (54,55). This effect is likely exacerbated during mitochondrial stress, such as rotenone-mediated inhibition of Complex I (56). The supplemental OA, LA, GLA, or ARA could also serve as a target for ROS-mediated peroxidation, which would cause membrane damage, further increasing ROS production and driving increased ESRE activation. We predict that substantial membrane damage may be sufficient to trigger ESRE activation, even if levels of MDT-15, FAT-6, or FAT-7 are low. This may also explain the importance of the lipid metabolism genes in ESRE activation. Polyunsaturated fatty acids (like LA, GLA, and ARA) are more commonly found in phosphoethanolamine, a major membrane lipid component, in the ER and mitochondria (57). Disruption of these genes is likely to perturb mitochondrial membrane lipid content, compromising ESRE signaling and mitochondrial health and stability.

Lipid peroxidation has other detrimental effects on cells. For example, lipid peroxidation induced by Complex I inhibition has been shown to cause ferroptosis (58). Ferroptosis is an iron-dependent form of programmed cell death whose role is recognized in an increasing number of diseases, including cancer, neurodegenerative diseases, and some autoimmune disorders (59). Interestingly, this may suggest a link between the ESRE response and the regulation of ferroptosis. One possibility is that the ESRE pathway limits ferroptotic activation by restoring cellular homeostasis before lipid peroxidation accumulates to levels sufficient to trigger cell death. An alternative possibility is that the ESRE pathway propagates a signal to help activate ferroptosis to limit further damage if mitophagy has failed. Investigation to test these possibilities is currently a priority.

Interestingly, one plausible mechanism for box C/D snoRNP complexes in the regulation of *fat-6* and *fat-7* has already been demonstrated to exist in humans. The box C/D snoRNA *SNORND88C* catalyzes the 2’-O-methylation of C3680 in 28S rRNA, increasing the translation of the *SCD1* mRNA (60). SCD1 increases the ratio of unsaturated to saturated fatty acids, limiting lipid peroxidation. There are several arguments that this mechanism may underlie our observations, suggesting that our observations may likewise extend to humans. First, the 28S rRNA sequence targeted for 2’-O-methylation in humans is conserved in *C. elegans*. Although a simple BLAST search failed to locate a homolog of *SNORD88C,* this may be a function of the different rates of mutation that are likely in the snoRNA compared to rRNA, which mutates at a considerably slower rate (61). Additionally, the C, C’, D, and D’ regions of the box C/D snoRNA, which are responsible for targeting the methylation, are both short and spread through the ∼100nt *SNORD88C* transcript, further complicating a simple sequence search. Second, we observed that *fib-1(RNAi)* mutants exhibited increased levels of *fat-7* mRNA but lower levels of FAT-7 protein upon mitochondrial stress than wild-type animals, which is consistent with observations by Wang et al. that 28S rRNA modification caused post-transcriptional regulations of SCD1, increasing its protein levels after methylation of the rRNA (60).

The ESRE pathway responds to mitochondrial damage, which can lead to autophagic degradation of mitochondria if it is unresolved. For this reason, we investigated whether there was a relationship between ESRE and the PINK-1/Parkin pathway. Using a PINK-1::GFP reporter, we looked at the effect of mitophagic activation under conditions of reduced (i.e., *mdt-15(RNAi)*, *fat-6(RNAi)*, or *fib-1(RNAi)*) or increased (i.e., fatty acid supplementation) ESRE activity. Our results revealed a novel regulatory activity for the ESRE pathway: reduced ESRE activity increased PINK-1::GFP accumulation, indicating that functional ESRE normally places a brake on mitophagic activation during mitochondrial stress. Interestingly, fatty acid supplementation under these conditions enhanced PINK-1::GFP accumulation, despite the limited function of the ESRE regulators. As noted above, this may be a result of increased lipid oxidation and causing further mitochondrial damage and mitophagy, despite increased ESRE activity.

Despite significant effort, we have so far been unable to identify a single transcription factor that is indispensable for ESRE gene transcription. Instead, multiple transcriptional regulators whose disruption partially reduces ESRE gene expression have been found, including the nematode-specific zinc-finger transcription factor SLR-2, the conserved ribosomal histidine hydroxyl transferase JMJC-1/NO66/RIOX1, the SWI/SNF nucleosome remodeling complex PBAF, the bZIP transcription factors ZIP-2, ZIP-4, CEBP-1, and CEBP-2, and the box C/D snoRNP machinery (9,26,33,62). The addition of MDT-15 to this list suggests that its known partners, such as NHR-49 (34), NHR-45 (63), HSF-1 (64), and SBP-1 (65) should be investigated as well. However, uncovering an unexpected relationship between MDT-15, box C/D snoRNPs, fatty acid desaturases, and the ESRE pathway helps provide insight into the complex regulation of mitochondrial homeostasis.

## Author contributions

Conceptualization N.V.K.; methodology and investigation L.A., A.C., E.T., M.W., and Y.A.; writing – original draft L.A.; review and editing, all authors; funding acquisition N.V.K.

## Materials and Methods

### C. elegans Strains

All *C. elegans* strains were maintained on nematode growth medium (NGM) plates seeded with *E. coli* strain OP50 as the food source and were maintained at 22°C (66), unless noted otherwise. *C. elegans* strains used in this study included N2 Bristol (wild-type) (67), SS104 (temperature-sensitive mutant that produces progeny at 15°C and is sterile at 25°C) |*glp-4(bn2)I*| (68), WY703 |*fdIs2* [*3XESRE*::GFP]; pFF4[*rol-6*(*su1006*)]| (62), BX106 |fat-6(tm331) IV| (69), BX153 |fat-7(wa36) V| (69), SJ4100 |*zcIs13* [*Phsp-6*::GFP]| (46), DMS303 |*nIs590* [*fat-7p*::FAT-7::GFP + *lin15*(+)] V| (36), PHX649 |*fat-6*(*syb649*[*fat-6p*::FAT-6::GFP]) IV| (37), COP262 |*knuSi221*[*fib-1p*::FIB-1(genomic)::eGFP::*fib-1* 3’ UTR + *unc-119*(+)]| (44), IJ1651|*yh44[mdt-15*::*degron*::*EmGFP]* (41), NVK90|*pink-1*(*tm1779*);houIs001{by*Ex655*[*Ppink-1*::PINK-1::GFP + *Pmyo-2*::mCherry]} (42), DA2123 |adIs2122 [lgg-1p::GFP::lgg-1 + rol-6(su1006)]| (44,45) and CL2166 |dvIs19 [(pAF15)gst-4p::GFP::NLS]| (49).

Synchronized worms were prepared by hypochlorite isolation of eggs from gravid adults, followed by hatching of the eggs in pure S Basal. 6,000 synchronized L1 larvae were reared on 10 cm NGM plates seeded with OP50. After transfer, worms were grown at 22°C for 48 hours before experiments, or three days for the next isolation of eggs. For synchronization of SS104 worms, 6,000 synchronized L1 larvae were reared on 10 cm NGM plates seeded with OP50 and grown at 15°C for five days for the next isolation of eggs. L4-stage hermaphrodite worms were used for all assays unless specified in the text.

### Bacterial strains

RNAi experiments in this study were done using RNAi-competent *E. coli* (HT115-based) obtained from the Ahringer or Vidal RNAi libraries (70,71). For experiments requiring OP50-based RNAi, plasmids isolated from RNAi-competent *E. coli* (HT115-based) were transformed into RNAi-competent *E. coli* (OP50). All strains were sequenced before use.

### RNA interference protocol

RNAi-expressing *E. coli* were cultured and seeded onto NGM plates supplemented with 25 μg/mL carbenicillin and 1 mM IPTG. For RNAi experiments starting at L1, 1500 synchronized L1 larvae were reared on 6 cm RNAi plates and grown at 22°C for 48 hours before imaging, exposure to chemical treatment, or pathogens. For double RNAi experiments, specifically pertaining to *Phsp-6*::GFP experiments, RNAi was mixed at a 3:1 ratio with *spg-7(RNAi)* or *EV* control, respectively.

### *C. elegans* liquid-based chemical exposure assays

Synchronized L4-stage worms were washed from NGM plates seeded with *E. coli* OP50, HT115-based RNAi-expressing *E. coli*, or OP50-based RNAi-expressing *E. coli* into a 15 mL conical tube and rinsed three times. For experiments involving the *3XESRE*::GFP strain, worms were sorted into a 96-well half-area plate (∼100 worms/well), and the total volume per well, including treatment, was 100 µL. Treatment with rotenone was done at a final concentration of 25 µM per well. Treatment with ascorbate was done at a final concentration of 25 mM per well. For liquid supplementation of fatty acids, a final concentration of 300 µM per well was used. For treatment with 1,10-phenanthroline, a final concentration of 4 mM per well was used. For treatment with CCCP, a final concentration of 78.2 µM per well was used. For treatment with nitrostyrene, a final concentration of 30 µM per well was used. DMSO was used as the solvent control for all of these treatments at a *v/v* ratio relative to the treatment.

For experiments involving the use of the FAT-7::GFP, FAT-6::GFP, or the PINK-1::GFP strains, worms were sorted into a 96-well full area plate (∼150 worms/well), and the total volume per well, including treatment, was 160 µL. Worms were imaged with a Cytation5 automated microscope every hour for eighteen hours at room temperature. Percent GFP area quantification was calculated by dividing the object sum area corresponding to “highly activated” GFP of worms in a well by the object sum area of all the worms within the well using Gen5 software. This was performed by setting up two “Cellular Analysis” calculations in Gen5; one calculates the object sum area of all the worms in a well by setting a low GFP channel threshold (highlights worms by their background fluorescence) and the other one calculates the object sum area of the “highly activated” GFP portions of worms in the well by setting a high GFP channel threshold.

### *C. elegans* agar-based fatty acid supplementation assays

Fatty acids were obtained from Nu-Chek-Prep, Inc. and were prepared as previously described, with the modification of using DMSO as the vehicle rather than NP-40, unless specified otherwise (72). Assays were performed by growing synchronized L1 worms on the indicated fatty acid or control media. Unless otherwise indicated, OA, LA, GLA, and ARA were used at a final concentration of 500 μM. Fatty acid supplementation assays recommend the addition of a surfactant, such as NP-40 (Tergitol), to the agar before autoclaving in order to increase the solubility of the fatty acid in the media. However, the fatty acid can also be delivered via DMSO as its vehicle. Same results were obtained in fatty acid supplementation experiments with the use of NP-40 and DMSO as the vehicle for fatty acid supplementation **(Fig. S1 D-F).**

### *C. elegans* liquid-based ROS Measurements

*C. elegans* ROS measurements were conducted as previously described, with some adjustments (73). Synchronized L4-stage *glp-4* worms were washed from 6 cm NGM plates seeded with OP50-based RNAi-expressing *E. coli* into a 1.5 mL tube and rinsed three times with S basal and Tween. Worms were sorted into a 96-well half-area plate (∼50 worms/well), and the total volume per well, including treatment, was 100 µL. To induce ROS, worms were treated with 78.2 µM CCCP, and to quantify/visualize ROS, worms were stained with a final concentration of 25 µM DCFDA per well. Worms were imaged, and fluorescence intensity was measured using a Cytation5 automated microscope at 0 hours, 4 hours, and 5 hours. Mean fluorescence intensity (MFI) was calculated by dividing the fluorescence intensity by the number of worms per well.

### Quantitative reverse transcriptase PCR (qRT-PCR)

30,000 worms were used for RNA extraction, and subsequent qRT-PCR was performed as previously described (74). Before RNA extraction, worms were reared on RNAi plates starting at the L1 stage. For treatment with rotenone, L4-stage worms were washed off plates, rinsed three times, and incubated for 8 hours in S Basal supplemented with 25 μM rotenone, 300 μM fatty acid + 25 μM rotenone, or a corresponding volume of DMSO control.

### Microscopy

For visualization of COP262 (FIB-1::GFP), IJ1651 (MDT-15::GFP), and DA2123 (LGG-1::GFP) worms were immobilized using 10 mM levamisole and transferred onto 3% noble agar slides. At least three biological replicates with 15 worms per replicate were imaged using a Zeiss ApoTome.2 Imager.M2, Carl Zeiss, Germany with 63x magnification. For COP262, hypodermic nuclei in the tail of each worm were imaged. For IJ1651, nuclei in the head were imaged. Area of fluorescent punctae was quantified using the surfaces tool on a 10.2.0 version of Imaris software. When using the Surfaces tool, machine learning-based segmentation was used to identify fluorescent punctae within a defined region of interest (ROI). Parameters were stored for batch to ensure each image was analyzed consistently across all replicates. For DA2123, fluorescent punctae in the terminal bulb of the pharynx were imaged. The terminal pharyngeal bulb was set as the ROI, and LGG-1 punctae were counted using the spots tool on a 10.2.0 version of Imaris software.

### Statistical Analyses

At least three biological replicates were performed for each experiment. RStudio (version 4.3.0) was used to perform statistical analyses. Student’s t-test was performed to calculate the significance of a treatment or condition when there were only two cohorts in the experimental setting. One-way analysis of variance (ANOVA) was performed to calculate the significance of a treatment or a condition when there were three or more cohorts in the experimental setting. Statistically significant results, as determined via ANOVA, were then followed by the post-hoc Dunnett’s test to calculate statistical significance or *p* values between each group compared to the control group. Dunnett’s test was indicated in graphs as follows: no star = not significant, \**p* < 0.05, ** *p* < 0.01, and *** *p* < 0.001.

### Motif enrichment and pattern matching

To identify regulatory sequences in the 5′ non-coding regions of *C. elegans* genes upregulated in response to ethanol stress (Class I and Class III genes only), LK56, and LK-*Pa*, 1 kilobase of sequence data upstream of the start codon of each gene was retrieved using the metazoan Regulatory Sequence Analysis Tools software (https://rsat.france-bioinformatique.fr/metazoa/).

Promoters were then searched for overrepresented motifs using the program MEME (Multiple Em for Motif Elicitation) (http://meme.sdsc.edu/meme/intro.html). This analysis was performed locally, using background models of the fourth order. Motif logograms were generated by MEME. Position weight matrix (shown below) was derived from MEME and used in subsequent pattern matching searches in RSAT.

|  |  |  |  |  |  |  |  |  |  |  |  |
| --- | --- | --- | --- | --- | --- | --- | --- | --- | --- | --- | --- |
| **A** | 0 | 0 | 0.999985 | 0 | 2.99999 | 2.000005 | 0 | 0.999985 | 0.999985 | 2.99999 | 7 |
| **C** | 9.000005 | 35 | 2.99999 | 6.000015 | 30.99999 | 7.999985 | 6.000015 | 32.00001 | 0 | 30.99999 | 4.999995 |
| **G** | 0 | 0 | 0 | 28.99999 | 0 | 21 | 0 | 0.999985 | 0 | 0 | 0 |
| **T** | 26 | 0 | 30.99999 | 0 | 0.999985 | 4.00001 | 28.99999 | 0.999985 | 34.00002 | 0.999985 | 23.00001 |

## Acknowledgements.

We thank Evan Park for assistance with FIB-1::GFP microscopy assays. Some strains were provided by the CGC, which is funded by NIH Office of Research Infrastructure Programs (P40 OD010440). This research was supported by NIH NIGMS R35GM129294 grant to NVK and a supplement for NVK’s NCI award R21CA280500 to LA (3R21CA280500-01A1S1) and training grant T32 AI055449-17 to LA, and Rice University fellowships to AC.

## Supporting Information Titles and Legends

**Figure S1. Oleic acid by itself does not trigger ESRE activation, and using NP-40 or DMSO as the vehicle for fatty acid supplementation elicits the same results. (A,D)** Fluorescent images of *3XESRE*::GFP worms reared on *EV, mdt-15, fat-6* or *fat-7* RNAis supplemented with OA and treated with DMSO **(A)** or rotenone **(D)**. **(B,E)** Curve representing ESRE activation by OA **(B)** or rotenone **(E)** across various hours. **(C,F)** Area under the curve quantification for ESRE activation by OA **(C)** or rotenone **(F)** across various hours. Supplementation of OA in panels **A-C** was delivered using DMSO as the vehicle. Supplementation of OA in panels **E-F** was delivered using NP-40 as the vehicle. In all panels, red stars indicate comparisons between fatty acid supplementation and no supplementation for each cohort. In all panels, black stars indicate comparisons between RNAi knockdowns and *EV(RNAi)* negative control in the non-supplemented cohort. Student’s t-test was performed to calculate the significance of a treatment or condition when there were two cohorts in the experimental setting. One-way analysis of variance (ANOVA) was performed to calculate the significance of a treatment or a condition when there were more than two cohorts in the experimental setting. No stars = not significant, *p < 0.05, ** p < 0.01, and *** p < 0.001.

**Figure S2. Supplementation of downstream PUFAs enhances ESRE activation. (A,D)** Fluorescent images of *3XESRE*::GFP worms reared on *EV*, *mdt-15,* or *fib-1* **(G)** RNAis supplemented with LA **(D),** GLA **(D)**, or ARA **(G)** and treated with 25 μM rotenone. **(B,E,H)** Curve representing ESRE activation by rotenone across various hours. **(C,F,I)** Area under the curve quantification for ESRE activation by rotenone across various hours. **(J)** Area of FIB-1 punctae in FIB-1::GFP worms reared on *EV* or *mdt-15* RNAis supplemented with ARA and treated with DMSO. In all panels, red stars indicate comparisons between fatty acid supplementation and no supplementation for each RNAi. In all panels, black stars indicate comparisons between RNAi knockdowns and *EV(RNAi)* negative control in the non-supplemented cohort. Student’s t-test was performed to calculate the significance of a treatment or condition when there were two cohorts in the experimental setting. One-way analysis of variance (ANOVA) was performed to calculate the significance of a treatment or a condition when there were more than two cohorts in the experimental setting. No stars = not significant, *p < 0.05, ** p < 0.01, and *** p < 0.001.

**Figure S3. Fatty acid supplementation enhances mitophagic but not UPR^mt^ activation. (A)** Fluorescent images of PINK-1::GFP worms reared on *EV, mdt-15, fat-6,* or *fib-1* RNAis supplemented with ARA and treated with 30 µM nitrostyrene or *v/v* DMSO.**(B,D,F)** Fluorescent images of *Phsp-6*::GFP worms reared on different RNAis supplemented with OA **(B)**, LA **(D)**, or ARA **(F)**. **(C,E)** Quantification of percent GFP area, which gives a readout of UPR^mt^ activation. In all panels, red stars indicate comparisons between fatty acid supplementation and no supplementation for each RNAi. In all panels, black stars indicate comparisons between RNAi knockdowns and *EV(RNAi)* negative control in the non-supplemented cohort. Student’s t-test was performed to calculate the significance of a treatment or condition when there were two cohorts in the experimental setting. One-way analysis of variance (ANOVA) was performed to calculate the significance of a treatment or a condition when there were more than two cohorts in the experimental setting. No stars = not significant, *p < 0.05, ** p < 0.01, and *** p < 0.001.

**Table S1. ESRE genes upregulated by exposure to 7% ethanol**

**Table S2. ESRE genes upregulated in response to liquid-based *P. aeruginosa* pathogenesis**

**Table S3. ESRE genes upregulated in response to LK56 treatment**

