## Supplemental Figures and Tables for "Box C/D snoRNPs and MDT-15/MED15 regulate mitochondrial surveillance and mitophagy via fatty acid metabolism"

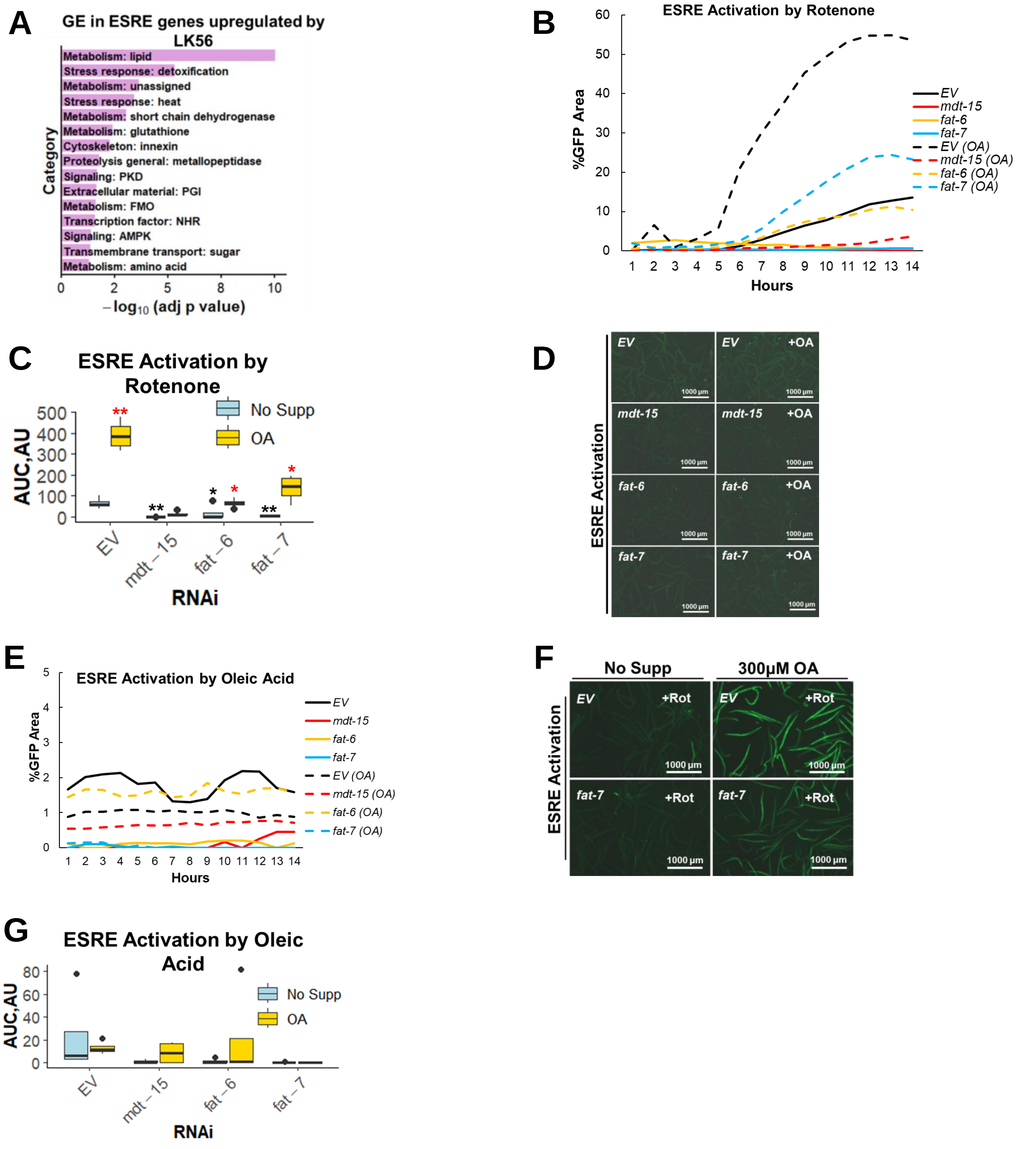
A
GE in ESRE genes upregulated by LK56
Category
Metabolism: lipid
Stress response: detoxification
Metabolism: unassigned
Stress response: heat
Metabolism: short chain dehydrogenase
Metabolism: glutathione
Cytoskeleton: innexin
Proteolysis general: metallopeptidase
Signaling: PKD
Extracellular material: PGI
Metabolism: FMO
Transcription factor: NHR
Signaling: AMPK
Transmembrane transport: sugar
Metabolism: amino acid
0 2 5 8 10
−log10 (adj p value)
B
ESRE Activation by Rotenone
%GFP Area
60 50 40 30 20 10 0
1 2 3 4 5 6 7 8 9 10 11 12 13 14
Hours
EV
mdt-15
fat-6
fat-7
EV (OA)
mdt-15 (OA)
fat-6 (OA)
fat-7 (OA)
C
ESRE Activation by Rotenone
AUC,AU
500 400 300 200 100 0
No Supp
OA
**
*
*
**
*
**
EV
mdt−15
fat−6
fat−7
RNAi
D
ESRE Activation
EV
EV +OA
mdt-15
mdt-15 +OA
fat-6
fat-6 +OA
fat-7
fat-7 +OA
1000 μm
E
ESRE Activation by Oleic Acid
%GFP Area
5 4 3 2 1 0
1 2 3 4 5 6 7 8 9 10 11 12 13 14
Hours
EV
mdt-15
fat-6
fat-7
EV (OA)
mdt-15 (OA)
fat-6 (OA)
fat-7 (OA)
F
No Supp
300μM OA
ESRE Activation
EV +Rot
EV +Rot
fat-7 +Rot
fat-7 +Rot
1000 μm
G
ESRE Activation by Oleic Acid
AUC,AU
80 60 40 20 0
No Supp
OA
EV
mdt−15
fat−6
fat−7
RNAi

**Figure S1**

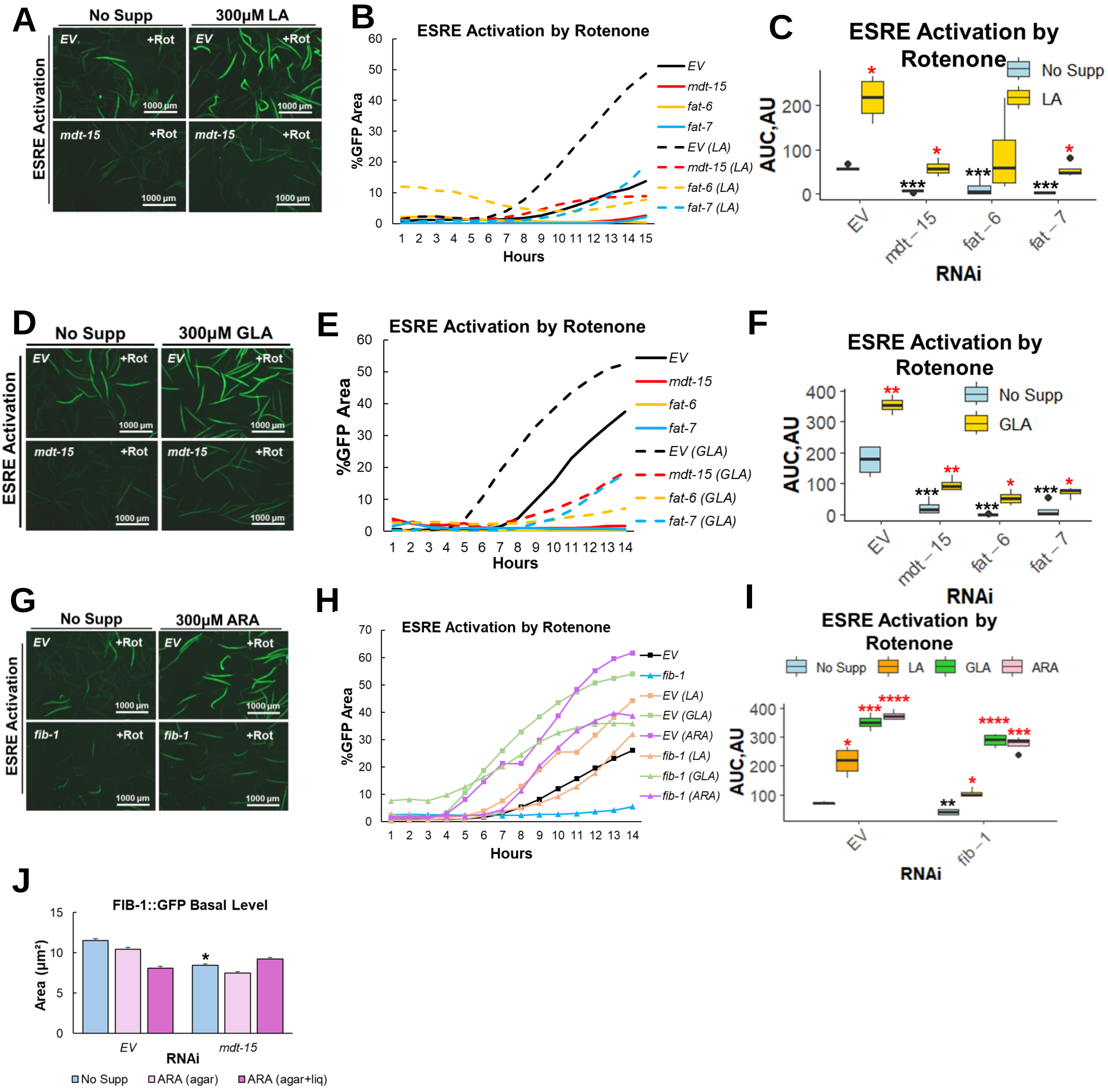
A
No Supp
300µM LA
ESRE Activation
EV
+Rot
mdt-15
1000 µm
B
ESRE Activation by Rotenone
%GFP Area
Hours
EV
mdt-15
fat-6
fat-7
EV (LA)
mdt-15 (LA)
fat-6 (LA)
fat-7 (LA)
C
ESRE Activation by Rotenone
No Supp
LA
AUC,AU
RNAi
EV
mdt−15
fat−6
fat−7
D
No Supp
300µM GLA
ESRE Activation
E
ESRE Activation by Rotenone
%GFP Area
Hours
EV (GLA)
mdt-15 (GLA)
fat-6 (GLA)
fat-7 (GLA)
F
ESRE Activation by Rotenone
No Supp
GLA
AUC,AU
RNAi
G
No Supp
300µM ARA
ESRE Activation
fib-1
H
ESRE Activation by Rotenone
%GFP Area
Hours
EV
fib-1
EV (LA)
EV (GLA)
EV (ARA)
fib-1 (LA)
fib-1 (GLA)
fib-1 (ARA)
I
ESRE Activation by Rotenone
No Supp
LA
GLA
ARA
AUC,AU
RNAi
EV
fib−1
J
FIB-1::GFP Basal Level
Area (µm²)
EV
mdt-15
RNAi
No Supp
ARA (agar)
ARA (agar+liq)

**Figure S2**

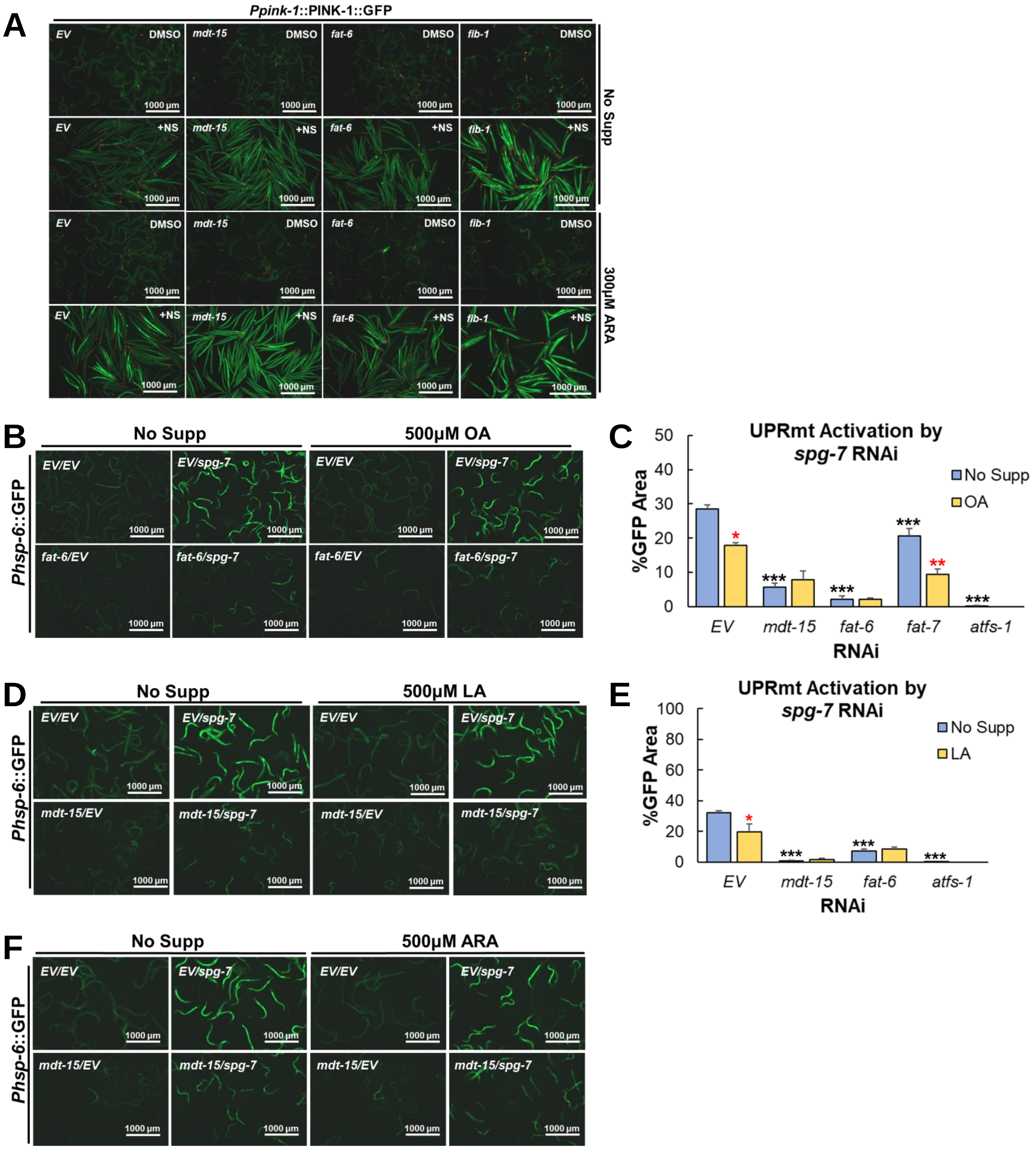
A
Ppink-1::PINK-1::GFP
EV DMSO
mdt-15 DMSO
fat-6 DMSO
fib-1 DMSO
EV +NS
mdt-15 +NS
fat-6 +NS
fib-1 +NS
No Supp
300µM ARA
1000 µm
B
No Supp
500µM OA
Phsp-6::GFP
EV/EV
EV/spg-7
fat-6/EV
fat-6/spg-7
C
UPRmt Activation by spg-7 RNAi
%GFP Area
No Supp
OA
EV
mdt-15
fat-6
fat-7
atfs-1
RNAi
*
**
***
D
No Supp
500µM LA
mdt-15/EV
mdt-15/spg-7
E
UPRmt Activation by spg-7 RNAi
LA
F
No Supp
500µM ARA

**Figure S3**

**Table S1. ESRE genes upregulated by exposure to 7% ethanol**

| Wormbase ID | Sequence ID | Gene Name |
|---|---|---|
| WBGene00002017 | T27E4.2 | *hsp-16.11* |
| WBGene00009824 | F47G4.3 | *gpdh-1* |
| WBGene00021610 | Y46H3D.6 | *nhr-237* |
| WBGene00015449 | C04F5.7 | *ugt-63* |
| WBGene00008592 | F08H9.4 | *F08H9.4* |
| WBGene00002026 | C12C8.1 | *hsp-70* |
| WBGene00009180 | F26H11.2 | *nurf-1* |

**Table S2. ESRE genes upregulated in response to liquid-based *P. aeruginosa* pathogenesis**

| Wormbase ID | Sequence ID | Gene Name |
|---|---|---|
| WBGene00001564 | C05E4.9 | *icl-1* |
| WBGene00044675 | Y119D3B.22 | *fbxa-76* |
| WBGene00003388 | F32A11.6 | *moe-3* |
| WBGene00011972 | T23G11.7 | *T23G11.7* |
| WBGene00016785 | C49G7.7 | *C49G7.7* |
| WBGene00012546 | Y37D8A.4 | *Y37D8A.4* |
| WBGene00006702 | Y71G12B.15 | *ubc-3* |
| WBGene00003655 | Y17D7A.3 | *nhr-65* |
| WBGene00003648 | R11G11.2 | *nhr-58* |
| WBGene00002016 | Y46H3A.3 | *hsp-16.2* |
| WBGene00022115 | Y71F9AL.10 | *Y71F9AL.10* |
| WBGene00002018 | Y46H3A.2 | *hsp-16.41* |
| WBGene00009328 | F32D8.3 | *F32D8.3* |
| WBGene00011300 | R107.5 | *R107.5* |
| WBGene00008476 | E03H4.8 | *E03H4.8* |
| WBGene00015248 | B0546.1 | *mai-2* |
| WBGene00007553 | C13G3.1 | *C13G3.1* |
| WBGene00002020 | T27E4.9 | *hsp-16.49* |
| WBGene00002019 | T27E4.3 | *hsp-16.48* |
| WBGene00002015 | T27E4.8 | *hsp-16.1* |
| WBGene00009692 | F44E5.5 | *F44E5.5* |
| WBGene00009691 | F44E5.4 | *F44E5.4* |
| WBGene00010625 | K07C5.2 | *K07C5.2* |
| WBGene00000514 | F22F7.5 | *ckb-4* |
| WBGene00012216 | W02D9.10 | *W02D9.10* |
| WBGene00003112 | F35G12.6 | *mab-21* |
| WBGene00019967 | R08F11.3 | *cyp-33C8* |
| WBGene00009588 | F40F11.3 | *F40F11.3* |
| WBGene00018164 | F38A5.7 | *sup-36* |
| WBGene00009534 | F38B7.3 | *F38B7.3* |
| WBGene00021977 | Y58A7A.3 | *Y58A7A.3* |
| WBGene00020596 | T20B5.3 | *oga-1* |

| | | |
|---|---|---|
| WBGene00017586 | F19B10.4 | *F19B10.4* |
| WBGene00010646 | K08C7.1 | *K08C7.1* |
| WBGene00011146 | R08D7.6 | *pde-2* |
| WBGene00000095 | C25A1.11 | *aha-1* |
| WBGene00015410 | C04A2.1 | *tbc-6* |
| WBGene00002026 | C12C8.1 | *hsp-70* |
| WBGene00021464 | Y39G10AR.6 | *ugt-31* |
| WBGene00012923 | Y47D3A.2 | *fbxa-128* |

**Table S3. ESRE genes upregulated in response to LK56 treatment**

| Wormbase ID | Sequence ID | Gene Name |
|---|---|---|
| WBGene00021927 | Y55F3AM.10 | *abhd-14* |
| WBGene00017874 | F28A10.6 | *acdh-9* |
| WBGene00000179 | ZK525.2 | *aqp-11* |
| WBGene00007605 | C15C8.3 | *asp-10* |
| WBGene00015163 | B0361.9 | *B0361.9* |
| WBGene00007254 | C01H6.4 | *C01H6.4* |
| WBGene00007599 | C15A11.7 | *C15A11.7* |
| WBGene00008032 | C39E9.8 | *C39E9.8* |
| WBGene00021491 | Y40B10A.6 | *comt-4* |
| WBGene00000830 | Y54G11A.6 | *ctl-1* |
| WBGene00019967 | R08F11.3 | *cyp-33C8* |
| WBGene00000982 | T11F9.11 | *dhs-19* |
| WBGene00000986 | R08H2.1 | *dhs-23* |
| WBGene00000967 | T02E1.5 | *dhs-3* |
| WBGene00017147 | EGAP2.1 | *EGAP2.1* |
| WBGene00044811 | F12E12.11 | *F12E12.11* |
| WBGene00017429 | F13D11.4 | *F13D11.4* |
| WBGene00017640 | F20D6.11 | *F20D6.11* |
| WBGene00009110 | F25D1.5 | *F25D1.5* |
| WBGene00018342 | F42A10.6 | *F42A10.6* |
| WBGene00009809 | F47B8.8 | *F47B8.8* |
| WBGene00018621 | F48G7.10 | *F48G7.10* |
| WBGene00018645 | F49F1.5 | *F49F1.5* |
| WBGene00010000 | F53F8.3 | *F53F8.3* |
| WBGene00010019 | F54B8.4 | *F54B8.4* |
| WBGene00018984 | F56F10.1 | *F56F10.1* |
| WBGene00010336 | F59F4.1 | *F59F4.1* |
| WBGene00001480 | H24K24.5 | *fmo-5* |
| WBGene00014030 | ZK637.13 | *glb-1* |
| WBGene00022196 | Y71H2B.7 | *gpa-17* |
| WBGene00009165 | F26E4.12 | *gpx-1* |
| WBGene00001755 | F11G11.2 | *gst-7* |
| WBGene00013073 | Y51A2D.4 | *hmit-1.1* |

| | | |
|---|---|---|
| WBGene00015471 | C05D9.2 | *lmp-2* |
| WBGene00021371 | Y37E11AR.4 | *nape-1* |
| WBGene00003609 | B0280.8 | *nhr-10* |
| WBGene00003724 | F44C8.2 | *nhr-134* |
| WBGene00003682 | Y41D4B.8 | *nhr-92* |
| WBGene00017671 | F21F3.1 | *pgal-1* |
| WBGene00018467 | F45E1.7 | *sdpn-1* |
| WBGene00012647 | Y39A1A.8 | *swt-4* |
| WBGene00011540 | T06E6.10 | *T06E6.10* |
| WBGene00011801 | T16G1.7 | *T16G1.7* |
| WBGene00020895 | T28D9.3 | *T28D9.3* |
| WBGene00007072 | AC3.7 | *ugt-1* |
| WBGene00010904 | M88.1 | *ugt-62* |
| WBGene00012445 | Y16B4A.2 | *Y16B4A.2* |
